# NET4 modulates the compactness of vacuoles in *Arabidopsis thaliana*

**DOI:** 10.1101/714774

**Authors:** Sabrina Kaiser, Ahmed Eisa, Jürgen Kleine-Vehn, David Scheuring

**Author notes:** equal contribution (Sabrina Kaiser, Ahmed Eisa). Correspondence should be addressed to David Scheuring.

## Abstract

The dimension of the plants largest organelle – the vacuole, plays a major role in defining cellular elongation rates. The morphology of the vacuole is controlled by the actin cytoskeleton but the mechanistic connection between them remains largely elusive. Recently, the NETWORKED (NET) family of membrane-associated, actin-binding proteins has been identified and represent potential candidates to impact on vacuolar morphology. Here, we show that NET4A localizes to highly constricted regions in the vacuolar membrane and contributes to the compactness of the vacuole. Using genetic interference, we found that deregulation of NET4 abundance impacts on vacuole morphogenesis and overexpression leads to more compact vacuoles. We moreover show that the NET4A-induced changes in vacuolar shape correlates with reduced cellular and organ growth in *Arabidopsis thaliana*. Our results demonstrate that NET4 modulates the compactness of vacuoles and reveal higher complexity in the regulation of actin-reliant vacuolar morphology.

## Results and Discussion

### NET4A shows a bead-on-a-string pattern at the tonoplast

Vacuolar size correlates and contributes to cell size determination, thereby defining cellular elongation rates (Löfke et al., 2015; Scheuring et al., 2016; Dünser et al., 2019). The actin cytoskeleton shows close proximity to the tonoplast and impacts on vacuolar morphology (Kutsuna et al., 2003; Scheuring et al., 2016). However, little is known about molecular players, mechanistically connecting the vacuolar membrane (tonoplast) with the actin cytoskeleton. The plant-specific NETWORKED (NET) family of membrane-associated, actin-binding proteins specifically link actin filaments to cell organelles (Deeks et al., 2012). The subfamily members NET4A and NET4B could impact on the actin-vacuole interface, because NET4A binds actin and overlaps with the tonoplast (Deeks et al., 2012).

To investigate the potential role of NET4 proteins in vacuolar morphology, we initially inspected NET4A∷NET4A-GFP localization in root epidermal cells of *Arabidopsis thaliana* seedlings. While NET4A-GFP is weakly detectable in meristematic cells (Löfke et al., 2015), we noted that it is preferentially expressed in the late meristematic zone, correlating with the onset of cellular elongation (Figure 1A). The root epidermis is regularly spaced between longer atrichoblast and shorter trichoblast cells, which later differentiate into non-hair and root-hair cells (Löfke et al., 2015). *NET4A* expression seems to precede in trichoblast cells and is more tightly associated with the onset of elongation in the atrichoblast cells (Figure 1A; Figure S1A-S1B). To confirm vacuolar localization of the NET4A-GFP signal, tonoplast staining using FM4-64 (Scheuring et al., 2015) was employed. NET4A-GFP did not only colocalize with the endocytic dye FM4-64 (Scheuring et al., 2015), but also displayed a more filamentous signal distribution in the cell cortex, suggesting a dual localization (Figure 1C). This cortical NET4A-GFP signal strongly resembles actin filaments, which is in agreement with its *in vitro* actin binding capacity (Deeks et al., 2012)

**Figure 1:**
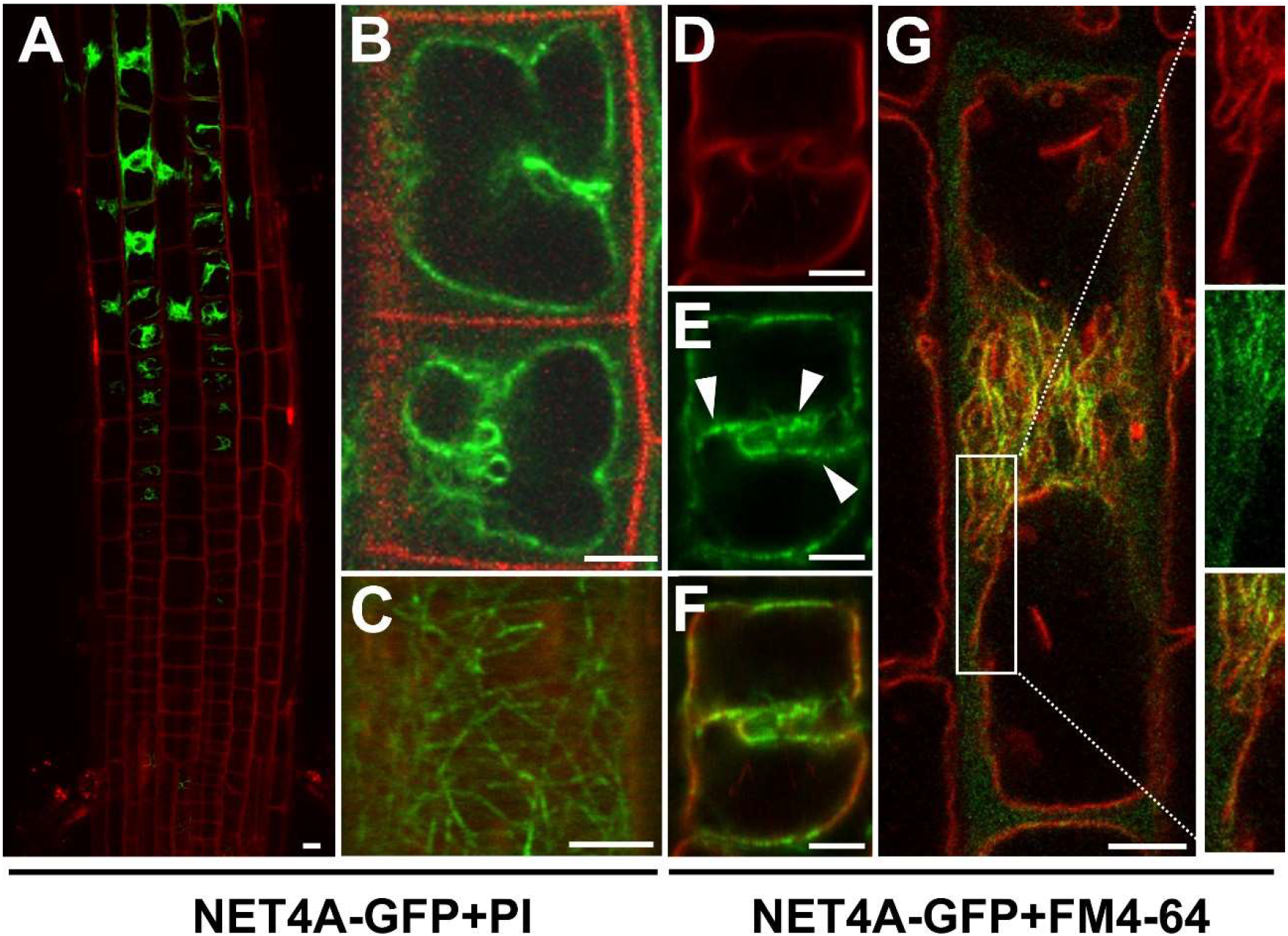
NET4A shows a bead-on-a-string pattern at the tonoplast. **(A)** NET4A-GFP expression under the endogenous promoter in Arabidopsis roots. Expression starts in the late meristem, initially only in trichoblast cells. **(B)** NET4A-GFP signal distribution in atrichoblast cells of the root epidermis. **(C)** Filamentous NET4A-GFP signals at the root cell cortex. **(D-F)** Vacuole staining by FM4-64 (3 h) shows NET4A localization preferentially at curved tonoplast areas **(G)** NET4A-GFP signal accumulation at constrictions in elongating root cells. Detail: areas with folded tonoplast membranes show stronger NET4A-GFP accumulation in comparison to expanding areas. White arrowheads highlight punctate signals. Propidium iodide was used to stain cell walls, FM4-64 to stain the tonoplast. Scale bars: 5 μm.

### NET4A localizes to highly constricted vacuolar membranes

NET4A-GFP distribution at the tonoplast was not uniform, but showed a stronger vacuolar label in the center of the cell, correlating with regions of higher vacuolar constrictions (Figure 1D-F). Vacuoles are highly constricted in meristematic cells and dramatically increase in volume during cellular elongation (Dünser et al., 2019). In agreement, in elongating cells the NET4A-GFP accumulation at the tonoplast correlated well with the remaining membrane constrictions (Figure 1G). In line with this, elongated root cells, possessing fully expanded vacuoles with no or only little constrictions, displayed only a very faint NET4A-GFP signal (Figure S1C).

To assess if NET4A-GFP accumulation indeed correlates with vacuolar constrictions, we next induced alterations in vacuolar morphology. The phytohormone auxin reduces vacuolar size by inducing smaller luminal vacuoles (Löfke et al., 2015; Scheuring et al., 2016). We used the synthetic auxin naphthalene acetic acid (NAA) and the auxin biosynthesis inhibitor kynurenine (kyn) to increase and decrease vacuolar constrictions in the root cells, respectively. In late meristematic cells, auxin treatment (200 nM NAA) induced vacuolar constrictions and a more uniform colocalization of NET4A-GFP with FM4-64 at the tonoplast (Figure 2A-C). On the contrary, kyn-induced reduction of vacuolar constrictions correlated with faint NET4A-GFP localization at the tonoplast (Figure 2A-C), being reminiscent of fully elongated cells. Auxin does not markedly interfere with NET4A-GFP intensity of already highly constricted vacuoles in meristematic cells (Löfke et al., 2015), but in late meristematic cells the overall signal intensity of NET4A was detectably increased and decreased in auxin treated and deprived cells, respectively (Figure 2D). To confirm this finding, we determined NET4A levels in response to increasing auxin concentrations by immunoblotting. Using a GFP antibody, we analyzed NET4A-GFP signal intensity in whole root extracts (Figure 2E). In a dosage-dependent manner, NAA application increased the NET4A-GFP protein amounts (Figure 2F). On the other hand, auxin application did neither elevate *NET4A* nor *NET4B* expression (Figure 2G), suggesting that auxin indirectly modulates the protein levels of NET4A.

**Figure 2:**
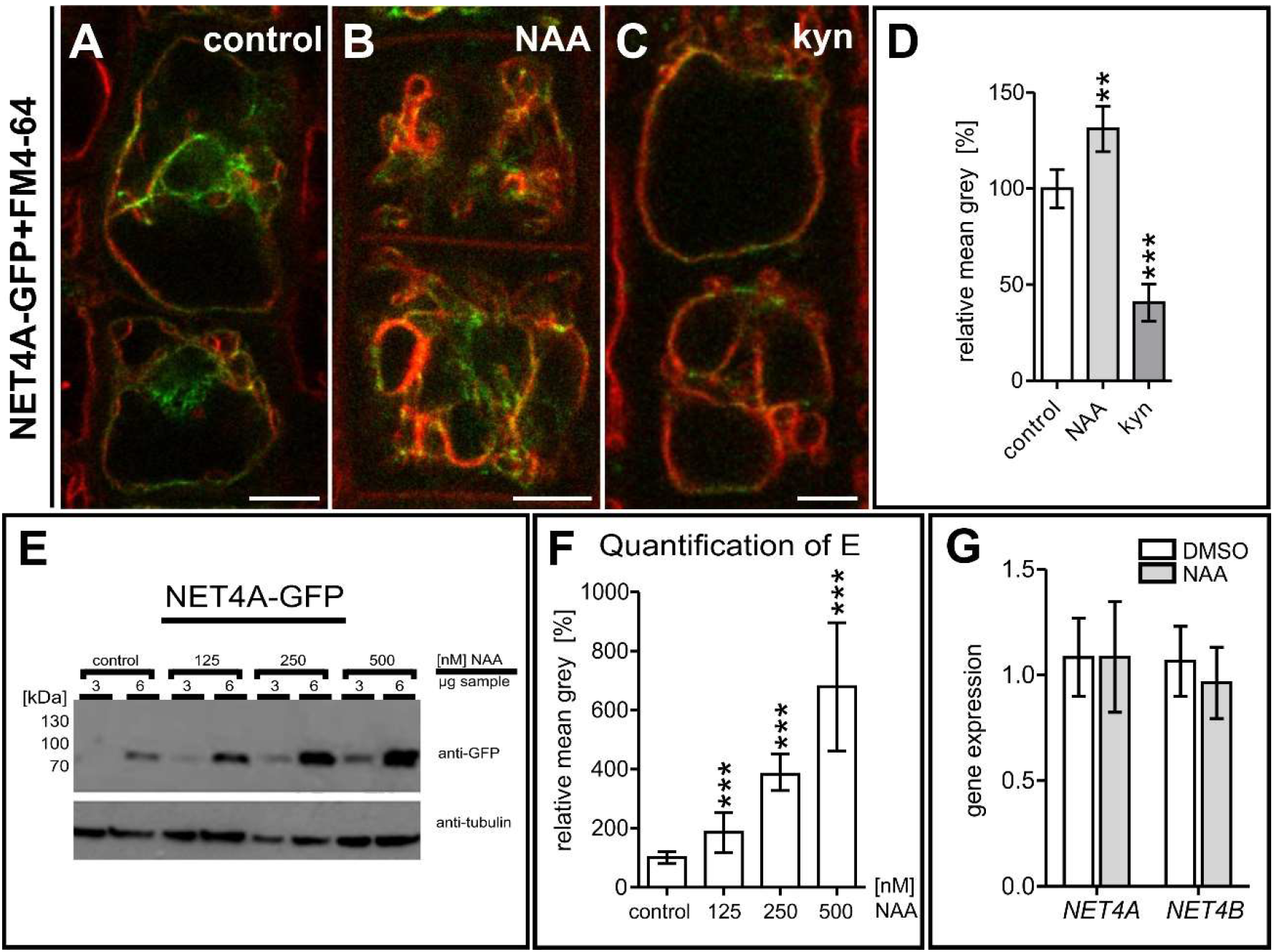
NET4A localizes to highly constricted vacuolar membranes. **(A-C)** Altered NET4A-GFP protein abundance upon exogenous auxin (NAA) application and depletion by the auxin biosynthesis inhibitor kynurenine (kyn). Seedlings were treated with DMSO as control (n=17), 250 nM NAA (n=15) and 3 μM kyn (n=15). Five cells per root were considered. **(D)** Quantification of signal intensity. **(E)** Western blot analysis of NET4A-GFP abundance upon rising NAA concentration. Arabidopsis root tissue was probed using a GFP-antibody. To avoid signal saturation, two different sample concentrations were loaded (3 and 6 μg). **(F)** Signal intensity of the individual NET4A-GFP bands were quantified in respect to the corresponding tubulin control bands. **(G)** To test for auxin-induced gene expression changes, qRT-PCR for both NET4 family members, *NET4A* and *NET4B*, was carried out. Error bars represent s.e.m. Student’s t-test, p-values: *p < 0.05; **p < 0.01; ***p < 0.001. Scale bars: 5 μm.

Our set of data suggests that NET4A is recruited to highly constricted regions of the tonoplast. This assumption is also in agreement with the fact that NET4 proteins contain a potential BAR-like domain, which are in general capable of sensing membrane curvature by binding preferentially to curved membranes (Peter et al., 2004).

### NET4A impacts on vacuolar morphology

To investigate NET4A function in maintaining vacuolar morphology, we isolated *net4a* and *net4b* knock-out lines (Figure S2A-D). We next investigated the NET4-dependent effect on vacuolar morphology in late meristematic cells, because they mark the onset of NET4A expression. We used the vacuolar morphology index (VMI) (Scheuring et al., 2016; Löfke et al., 2015; Dünser et al., 2019), which depicts the size of the biggest luminal vacuolar structure and is highly sensitive to reveal alterations in vacuolar morphology. *net4a-1* (SALK_017623) and *net4b* (SALK_056957) mutants both displayed more spherical vacuoles accompanied with increased VMI (Figure S3E-H). This suggests that NET4 association at the constricted vacuolar membranes impact on vacuolar morphology.

On the other hand, the *net4a net4b* double mutant (Figure S2I) also showed more spherical vacuoles, but the VMI was largely not distinguishable from the single mutants (Figure 3B, E; Figure S3F and S3H), proposing higher redundancy in the actin-vacuole pathway.

Next we generated 35S∷NET4A-GFP (NET4A-GFP^OE^) overexpression lines to assess ectopic NET4A expression. NET4A-GFP^OE^ lines showed expression throughout the root, including the meristematic region. The subcellular characteristics of NET4A-GFP^OE^ remained, showing the filamentous signal at the cell cortex and the enhanced labelling of constricted tonoplast membranes (Figure S3A-3F). Interestingly, trichoblast cells, possessing more compact and constricted vacuoles, showed a higher signal intensity when compared to atrichoblast cells (Figure SG-3I). To assess the effect of NET4A overexpression on vacuolar morphology, the VMI for NET4A-GFP^OE^ was determined. Markedly, NET4A overexpression lines showed more roundish vacuoles (Figure 3C), and increased VMI in comparison to the control (Figure 3D). Accordingly, we conclude that NET4A overexpression as well as *net4a* loss-of-function induces more roundish vacuoles. This finding is reminiscent to the stabilization and depolymerization of the actin cytoskeleton, which also both induce more roundish vacuoles and increase the VMI (Scheuring et al., 2016).

**Figure 3:**
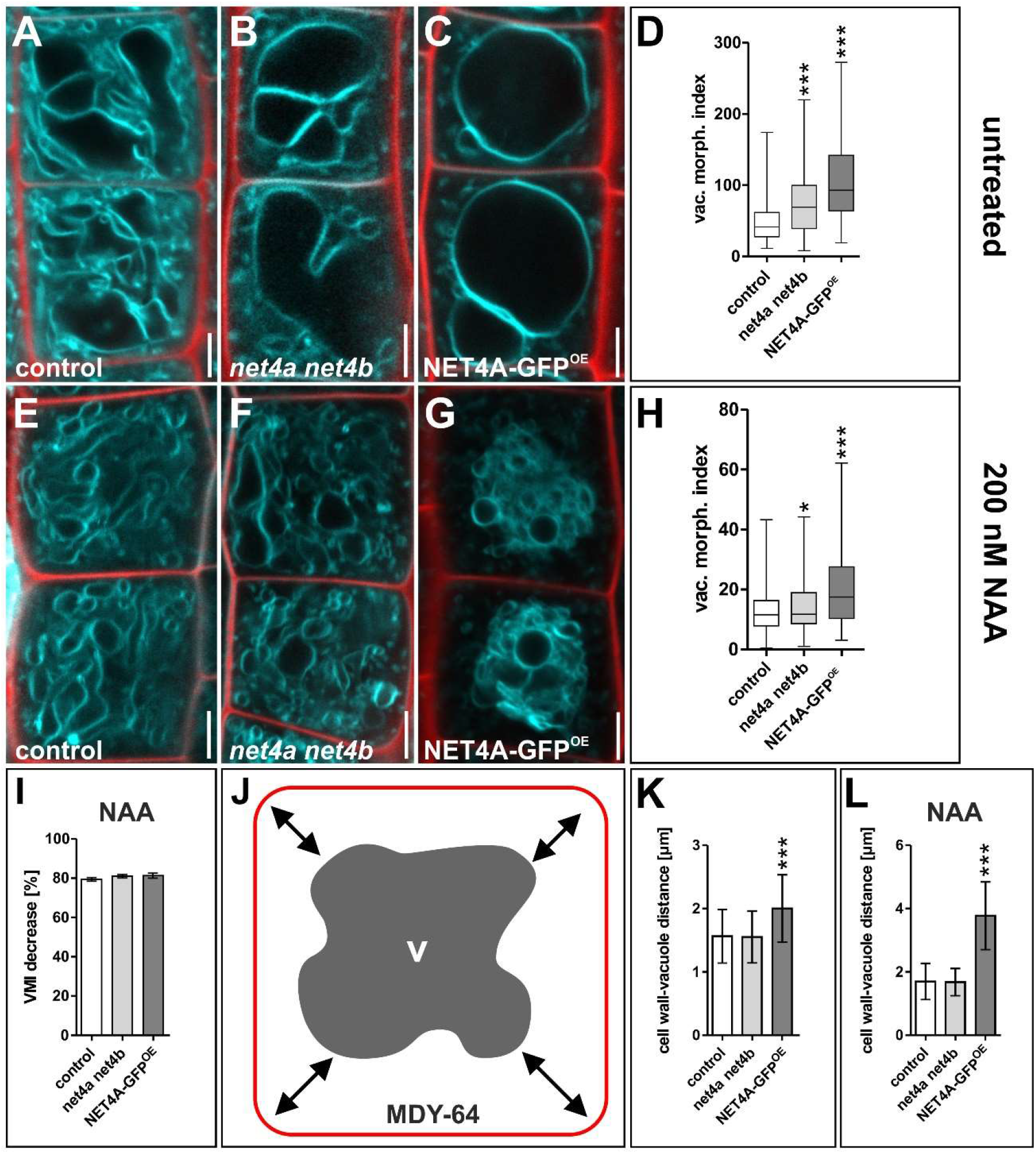
NET4 abundance impacts on vacuolar morphology, modulating compactness. **(A-D)** Vacuolar morphology in the late root meristem of the *net4a net4b* double knockout line (n=145) and of the NET4A-GFP line driven by the 35S promotor (n=99) in comparison to the Col-0 control (n=334). **(E-H)** Vacuolar morphology upon auxin treatment (200 nM NAA) in the control (n=147), net4a ne4b (n=153) and NET4A-GFP^OE^ (n=88) line. **(I)** Relative VMI decrease. **(J)** Vacuolar compactness was assessed by measuring plasma membrane to vacuole distance based on MDY-64 staining. The distance was measured in every cell corner and the mean calculated. **(K)** Quantification of plasma membrane to vacuole distance in *net4a net4b* (n=72) and NET4A-GFP^OE^ (n=22) in comparison to the Col-0 control (n=75). **(L)**. Quantification of auxin-induced changes (200 mM, 20h) of plasma membrane-vacuole distance in the control (n=59), *net4a net4b* (n=42) and NET4A-GFP^OE^ (n=20). Columns of bar charts represent mean values, error bars represent s.e.m. Box limits of boxblots represent 25th percentile and 75th percentile, horizontal line represents median. Whiskers display min. to max. values. Student’s t-test, p-values: *p < 0.05; **p < 0.01; ***p < 0.001. Scale bars: 5 μm.

### NET4 and auxin spatially define vacuolar occupation within the cell

Next, we used auxin-induced vacuolar constrictions, to further assess the contribution of NET4 to controlling vacuolar morphology. Following the exogenous application of auxin (200 nM, 20h), vacuoles of *net4a net4b* double mutants and the NET4A-GFP^OE^ line were slightly constricted when compared to wild type seedlings (Figure 3E-3J). However, considering that untreated *net4a net4b* double mutants and the NET4A-GFP^OE^ lines showed already more roundish vacuoles and higher VMIs, the relative responses were not distinguishable from wild type seedlings (Figure 3I).

Based on our data, we conclude that NET4 activity affects vacuolar morphology, but does not have a major impact on the auxin effect on vacuolar shape.

However, when inspecting vacuoles of NET4A-GFP^OE^ we noted that vacuoles seemed more condensed around the nucleus, showing a larger distance to the plasma membrane (Figure 3J). Accordingly, we quantified vacuolar distance to the plasma membrane in the respective genotypes and revealed that NET4A-GFP^OE^ indeed affected this cellular trait (Figure 3K). Auxin treatments did not alter the plasma membrane to vacuole distance in wild type, but considerably increased cell wall to vacuole distance in NET4A-GFP^OE^ (Figure 3L). Based on this set of data, we conclude that NET4A overexpression induces compactness of the vacuole.

### NET4A-dependent compacting of the vacuole correlates with reduced cell size and root organ growth

Changes in vacuolar morphology correlate with stomata movement (Andrés et al., 2014) and overall cellular elongation rates (Hawes et al., 2001; Singh et al., 2014; Löfke et al., 2015; Dünser et al., 2019). Therefore, we initially investigated whether NET4 proteins impact on cell size in the later meristematic region (Figure 4A; (Barbez et al., 2017)). While cell size in the *net4a net4b* double mutant remained unchanged in comparison to the control, NET4A-GFP^OE^ showed a significant reduction in cell length (Figure 4B). In agreement, the altered cell size determination correlated with reduced root organ growth (Figure 4C-E).

**Figure 4:**
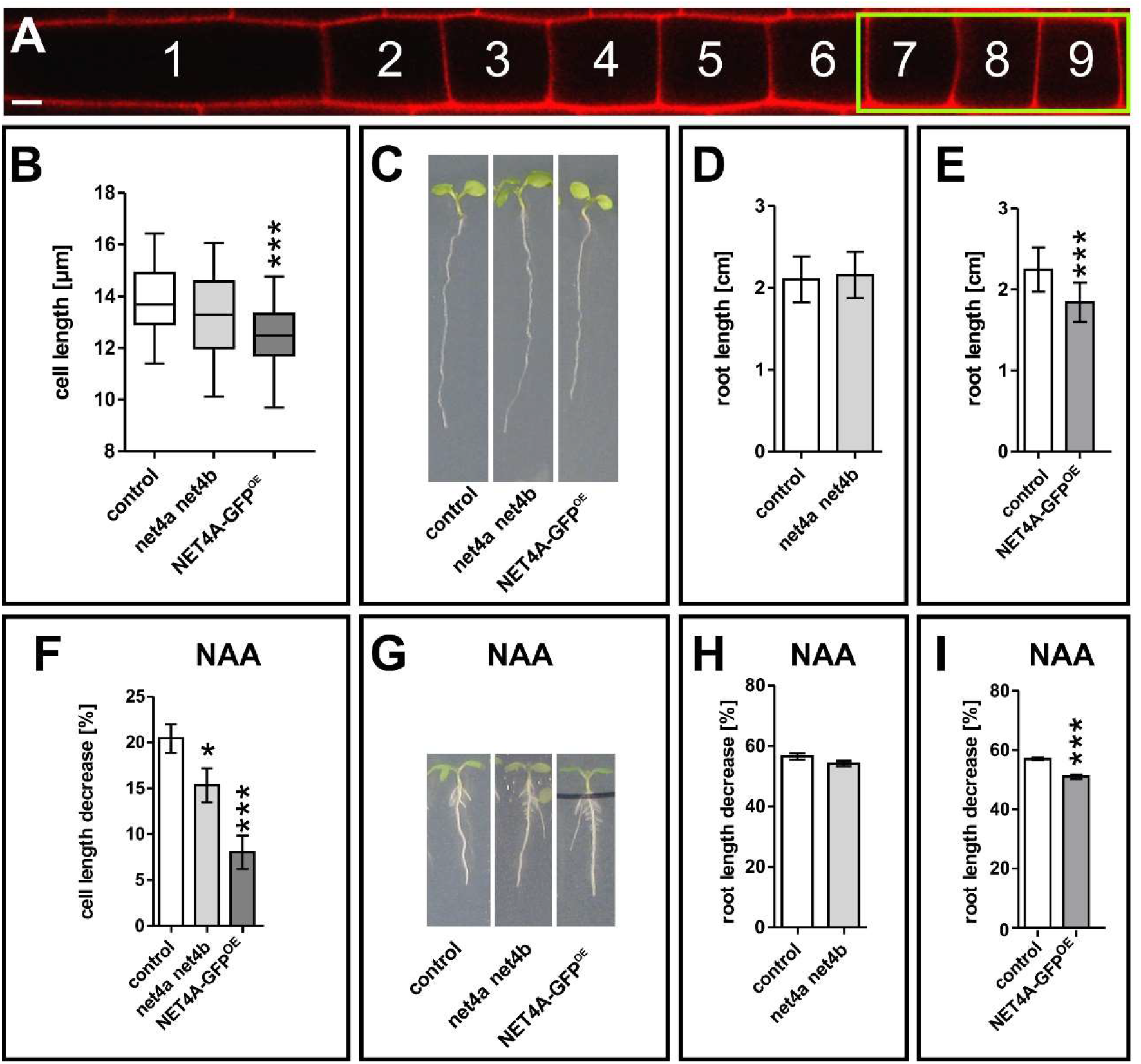
NET4A-dependent compacting of the vacuole correlates with reduced cell size and root organ growth. **(A)** Representative atrichoblast cell file for cell length quantification. The green box marks the three quantified cells per file (7 to 9). **(B)** Cell length of the *net4a net4b* (n=81) double knockout and the NET4A-GFP^OE^ (n=72) lines in respect to the control (n=147). **(C-E)** root length of the *net4a net4b* (n=27) double knockout and the NET4A-GFP^OE^ (n=24) lines in respect to the control (n=52). One representative experiment is shown. **(F)** Auxin-dependent (200 nM NAA, 20 h) cell length decrease of the double knockout (n=81) and NET4A-GFP^OE^ (n=81) in comparison to the control (n=147). **(G)** Root length of the control (n=50), *net4a net4b* (n=29) and NET4A-GFP^OE^ (n=26) upon auxin treatment (125 nM NAA, 6 d). One representative experiment is shown. **(H and I)** Quantification of auxin-dependent root length decrease. Columns of bar charts represent mean values, error bars display s.e.m. Box limits of boxblots represent 25th percentile and 75th percentile, horizontal line represents median. Whiskers display min. to max. values. Student’s t-test, p-values: *p < 0.05; **p < 0.01; ***p < 0.001. Scale bar: 5 μm.

Considering that auxin treatment was additive to the NET4A effect on the compactness of vacuoles, we next assessed cell length and root length changes upon NAA treatment in the *net4a net4b* mutant and in the NET4A-GFP^OE^ lines. The atrichoblast cell length of *net4a net4b* as well as NET4A-GFP^OE^ were partially resistant to auxin (200 nM NAA) when compared to wild type (Figure 4F). NET4A overexpression also caused a reduced root growth sensitivity to exogenous auxin (125 nM NAA) (Figure 4G-4I).

Taken together, we show that NET4A localizes to highly constricted regions in the vacuolar membrane and contributes to the compactness of the vacuole. We moreover show that the NET4A-induced changes in vacuolar shape also impacts on cellular and organ growth.

## Concluding Remarks

The plant vacuole is essential for development and growth (Schumacher et al., 1999; Rojo et al., 2001). In growing cells, vacuole size and cell size correlates (Owens and Poole, 1979; Berger et al., 1998; Löfke et al., 2013). Thus, a role of the vacuole in cell size determination has been proposed (Löfke et al., 2015), which was recently supported by the finding that increased vacuolar volume allows for rapid cellular elongation (Scheuring et al., 2016; Krüger and Schumacher, 2018; Dünser et al., 2019). However, the underlying forces, regulating vacuolar morphology, are not yet well understood. Previously, it was shown that the structural organization and dynamics of the vacuole relies on the interaction between cytoskeleton and the tonoplast (Higaki et al., 2006; Scheuring et al., 2016). It has been suggested that the actin cytoskeleton plays a major role during vacuole inflation in the late meristematic region (or transition zone) shortly prior the onset of cellular elongation. Here, it could provide the force to bring vacuolar structures into close proximity to allow for homotypic fusion events. In yeast, it has been reported that actin is enriched at the vacuole at sites where fusion occurs (Eitzen et al., 2002). This involves tethering, docking and fusion processes which in turn are dependent on Rab-family GTPases, vacuolar SNAREs and homotypic fusion and vacuole protein sorting (HOPS) complex (Zhang et al., 2014). In plants, pharmacological actin interference using profilin led to the disappearance of transvacuolar strands, ceased cytoplasmatic streaming and thereby affects vacuolar shape as well. (Staiger et al., 1994). Furthermore, actin interference has been shown to inhibit vacuole fusion during stomatal opening (Li et al., 2013). The role of NET4 within the regulation of vacuolar morphology was characterized due to its actin binding capacity (Deeks et al., 2012) and localization at the tonoplast (Figure 1). We could show that NET4 abundance is crucial to maintain vacuolar morphology and impacts subsequently on cell size and root length. However, compared to the importance of the actin myosin system (Scheuring et al., 2016), the loss of *NET4A* and *NET4B* has only mild impacts on vacuolar morphology and cell size. Accordingly, we conclude that higher molecular complexity provides a high level of redundancy for tethering the actin cytoskeleton to the vacuolar membranes. In metazoans, a variety of adaptor protein (e.g. spectrin, filamin) are known to provide specific contact sites for the actin cytoskeleton and various membranes (Wang et al., 2014). However, most of these protein families are not present in plants and it needs to be seen how these molecular functions are achieved in plants.

## Material and methods

### NET4 gene accession codes

Sequence data from this article can be found in The Arabidopsis Information Resource (TAIR; http://www.arabidopsis.org/) or GenBank/EMBL databases under the following accession numbers: NET4A (At5g58320) and NET4B (At2g30500).

### Plant Material, Growth Conditions and DNA Constructs

*Arabidopsis thaliana*, *Columbia 0* (*Col-0*) ecotype was used as control. The transgenic line NET4A∷NET4A-GFP has been described previously (Deeks et al., 2012). 35S∷NET4A-GFP was constructed using Gateway cloning. The NET4A coding sequence was amplified via PCR using root cDNA from 8 days old Arabidopsis seedlings. The primers are listed in table 1. The cDNA fragment was cloned into the pDONR221 (Invitrogen) using BP-clonase according to the manufacturer’s instructions. Then, the coding sequence was transferred from the entry vector into the destination vector pH7WG2 (Karimi et al., 2005) using the LR clonase from Invitrogen. Transformation into Arabidopsis thaliana Col-0, using the floral-dip method was carried out as described before (Barbez et al., 2012).

The insertion lines *net4a-1 (SALK_017623)* and *net4b (SALK_056957)* were obtained from the Nottingham Arabidopsis Stock Centre (NASC). *net4a net4b* was generated by crossing *net4a-1* and *net4b*. For identification of homozygous lines, genotyping of all insertion lines was performed using the NASC-recommended primers (table 1). Insertion sites were located by Sequencing within the 3th Exon (*net4a-1*) and the promoter (*net4b*), respectively. Gene knockout of the respective NET4 transcript was shown by qRT-PCR.

Seeds were surface sterilized with ethanol, plated on solid ½ Murashige and Skoog (MS) medium, pH 5.7-5.8 (Duchefa), containing 1 % (w/v) sucrose (Roth), 2.56 mM MES (Biomol) and 1 % (w/v) Phytoagar (Duchefa), and stratified at 4 °C for 1-2 days in the dark. Seedlings were grown in vertical orientation at 20-22 °C under long day conditions (16 h light/8 h dark).

### Chemicals and Treatments

α-naphthaleneacetic acid (NAA) was purchased from Duchefa; L-kynurenine (Kyn) from Sigma; and BCECF-AM, FM4-64, MDY-64 and propidium iodide (PI) from Life Technologies. All chemicals except PI were dissolved in dimethyl sulfoxide (DMSO). NAA and Kyn were applied in solid ½ MS medium, the dyes BCECF-AM, FM4-64 and MDY-64 in liquid ½ MS medium, and PI in distilled water.

### RNA Extraction and Quantitative Real Time PCR (qRT-PCR)

Total RNA of seedlings was extracted using either the innuPREP Plant RNA kit (analytic-jena) or the NucleoSpin RNA Plant kit (Macherey-Nagel). If necessary, RNA samples were additionally treated with DNase I (New England Biolabs) and purified using the RNA Clean and Concentrator kit (Zymo Research). DNA-free RNA samples were reverse transcribed using the iScript cDNA synthesis kit (Bio-Rad). All steps were performed according to the manufacturers recommendations. qPCR was performed with the iQ SYBR Green Supermix (Bio-Rad) or (in case of *net4a net4b* analysis) the PerfeCTa SYBR Green SuperMix for iQ (Quanta BioSciences) in a C1000 Touch Thermal Cycler equipped with a CFX96 Touch Real-Time PCR Detection System (Bio-Rad). The reaction procedure was set to a 95 °C heating step for 3 min, followed by 40 cycles of denaturing at 95 °C for 10 s and either separated annealing at 55 °C for 30 s and elongation at 72 °C for 30 s or (in case of *net4a net4b* analysis) combined annealing-extension at 60 °C for 40 s. Target specific and control primers used for quantification were given in table 1. Expression levels were normalized to the expression levels of either Ubiquitin 5 (UBQ5) or (in case of *net4a net4b* analysis) Actin 2 (ACT2). Relative expression ratios were calculated according to the so-called “delta-delta C_t_” method (Livak and Schmittgen, 2001) or the efficiency corrected calculation model (Pfaffl et al., 2004). Statistical analyzes were performed using Excel or REST 2009 software (www.gene-quantification.info).

### Phenotype Analysis

The quantification of vacuolar morphology, compactness and occupancy as well as cell length was carried out using 6-7-days-old seedlings. To evaluate changes upon kynurenine or auxin treatment, seedlings were transferred to solid ½ MS medium supplemented with 2 μM Kyn or 200 nm NAA 18-22h prior to image acquisition. For the analysis of the vacuolar morphology index, confocal sections above the nucleus of the root epidermis were acquired (Löfke et al., 2015). Calculations of the vacuolar morphology index were carried out in 4 late (distal) meristematic cells of atrichoblast files, respectively, as described previously (Dünser et al., 2019). For the quantification of plasma membrane – vacuole distance, the same cells of the late meristematic region were analyzed. Therefore, the distance from the plasma membrane corner to the first tonoplast structure in diagonal reach was measured for all four corners of a cell, then, mean was calculated and considered as single value for the plasma membrane – vacuole distance per cell. Cell length was measured in 3 early (proximal) meristematic cells of atrichoblast cell files (Figure 4A) as described before (Barbez et al., 2017). For measurements of signal intensity (mean gray value) a defined detector gain was used and individual cells in the late meristematic zone of the root were quantified. All confocal images taken to assess the vacuolar morphology index, plasma membrane–vacuole distance and cell length were analyzed using Fiji software. Z-stack confocal images taken to assess the vacuolar occupancy of the cell were further processed using Imaris software. For root growth determination, 6-7- days-old seedlings grown vertically on ½ MS medium plates were used and root length quantified using Fiji software. To analyze auxin dependent changes in root growth, solid ½ MS medium supplemented with 125 nm NAA was used. Statistical evaluation for all data were performed using Graphpad Prism 5 software.

### Identification of the BAR-like domain region

In order to predict conserved domains on the open reading frames of NET4A and NET4B, the protein sequences were retrieved from protein database at National Center for Biotechnology Information (NCBI: https://www.ncbi.nlm.nih.gov/) in FASTA format on 23 July 2019. In order to check for the functional units and patterns of these sequences, ExPASy PROSITE was used (http://prosite.expasy.org) (Sigrist et al., 2010). Here we identified the N-terminal actin-binding domain (NAB) stretching between amino acid position 21 and 101 for both NET4A and NET4B. In parallel, we also performed a conserved domain search using NCBI Conserved Domain Search (https://www.ncbi.nlm.nih.gov/Structure/cdd/wrpsb.cgi). Here, we identified an N-terminal localized kinase interacting protein 1 (KIP1) domain and a C-terminal localized structural maintenance of chromosomes (SMC) domain for both isoforms. No N-terminal actin-binding domain (NAB) was identified in either isoform in this search. We used the Simple Modular Architecture Research Tool (SMART) to search for regions of the full-length proteins that were deemed irregular (http://smart.embl-heidelberg.de/) (Schultz et al., 1998). Indeed, an unknown amino acids and low complexity sequence was identified stretching between amino acid positions 113 and 193, and 107 and 155, for NET4A and NET4B, respectively. Both sequence stretches start at positions directly after the predicted NAB domain. To identify domain architectural homology in the unknown protein sequences, both full-lengths and the first 200 amino acids sequences were used in the Conserved Domain Architecture Retrieval Tool (CDART) (https://www.ncbi.nlm.nih.gov/Structure/lexington/lexington.cgi?cmd=rps/) repository search (Geer et al., 2002). All results were extracted and filtered according to membrane proteins containing either an N-terminal KIP domain and/or C-terminal SMC domain. Several hits displayed proteins containing BAR domains.

### Western Blotting

Roots of 7-days-old seedlings expressing NET4A∷NET4A-GFP were homogenized in liquid nitrogen and solubilized in extraction buffer (200 mM Tris pH 6.8, 400 mM DTT, 8 % SDS, 40 % glycerol, 0.05 % bromphenol blue). After incubation at 95 °C for 5-10 min, samples were centrifuged and the proteins contained in the supernatant separated using SDS-PAGE (10 % gel). For blotting a polyvinylidene difluoride (PVDF) membrane (Immobilon-P, pore size 0.45 μm, Millipore) was used and after blocking with 5 % skim milk powder in TBST (150 mM NaCl, 10 mM Tris/HCl pH 8.0, 0.1 % Tween 20), the membrane was probed with a 1:20000 dilution of mouse anti-GFP antibody (JL-8, Roche) or mouse anti-alpha-tubulin antibody (B-511, Sigma). As secondary antibody, a 1:20000 dilution of horseradish peroxidase-conjugated goat anti-mouse antibody (pAB, Dianova) was used. Signals were detected using the SuperSignal West Pico chemiluminescent substrate detection reagent (Thermo Scientific) and quantified using Fiji software. Signal intensities of GFP were normalized to alpha-tubulin and statistical evaluation was performed using Graphpad Prism 5 software.

### Confocal Microscopy

For live cell imaging, except FM4-64 staining was performed, roots were mounted in PI solution (0.01 mg/ml) to counterstain cell walls. FM4-64 and MDY-64 staining of the tonoplast was performed as described before (Scheuring et al., 2015). For 3D imaging, vacuoles were stained with BCECF-AM (10 μM solution in ½ MS liquid medium) for at least 1.5 h in the dark (Scheuring et al., 2015). For staining of auxin treated samples, the staining solution was supplemented with NAA in the respective concentration. Confocal images were acquired using either a Leica SP5 (DM6000 CS), TCS acousto-optical beam splitter confocal laser-scanning microscope, equipped with a Leica HC PL APO CS 20 × 0.70 IMM UV objective and Leica HCX PL APO CS 63 × 1.20. water-immersion objective or a Zeiss LSM880, AxioObserver Z1 confocal laser-scanning microscope, equipped with a Zeiss C-Apochromat 40x/1.2 W AutoCorr M27 water-immersion objective. Fluorescence signals of MDY-64 (Leica-System: excitation/emission 458 nm/465-550 nm, Zeiss-System: excitation/emission 458 nm/473- 527 nm), GFP and BCECF (Leica: excitation/emission 488 nm/500-550 nm, Zeiss: excitation/emission 488 nm/500-571 nm), FM4-64 and PI (Leica-System: excitation/emission 561 nm/599-680 nm (FM4-64), 644-753 nm (PI), Zeiss-System: excitation/emission 543 nm/580-718 nm (PI)) were processed using Leica software LAS AF 3.1, Zeiss software ZEN 2.3 (black and blue edition) or Fiji software. For double labeling, images were acquired using sequential scan mode to avoid channel crosstalk. Z-stacks were recorded with a step size of 540 nm, resulting in approximately 25-35 single images.

## Acknowledgments

We would like to thank Patrick Hussey for sharing published material, Kai Dünser and Christian Löfke for technical help and Matthias Hahn for critical reading of the manuscript. This work was supported by the DFG (SCHE 1836/4-1).

