## Supplementary material for "NET4 modulates the compactness of vacuoles in *Arabidopsis thaliana*": Figure S1

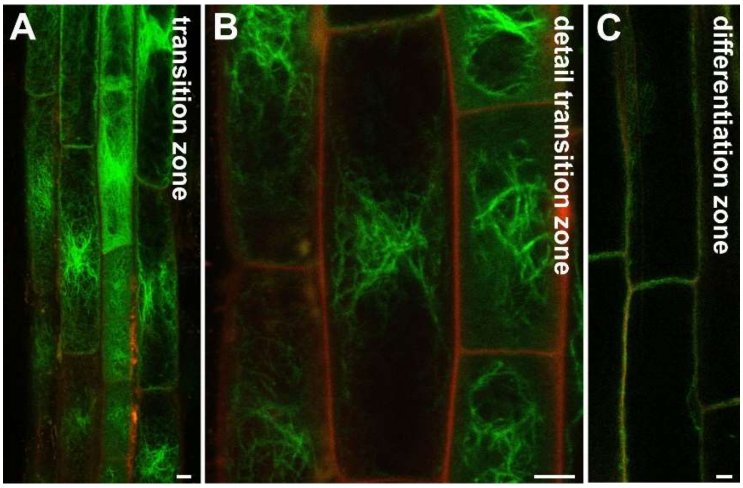


**Figure S1: Spatial localization of NET4A.** NET4A-GFP expression under its endogenous promoter in Arabidopsis roots. Expression starts in the late meristem, initially only in trichoblast cells (Figure 1A). **(A and B)** In (larger) atrichoblast cells, expression starts in the transition zone and continuous until the elongation zone. **(C)** NET4A-GFP is only weakly expressed in the differentiation zone. Propidium iodide (red) was used to stain cell walls. Scale bars: 5 µm.
