## Supplementary material for "NET4 modulates the compactness of vacuoles in *Arabidopsis thaliana*": Figure S2

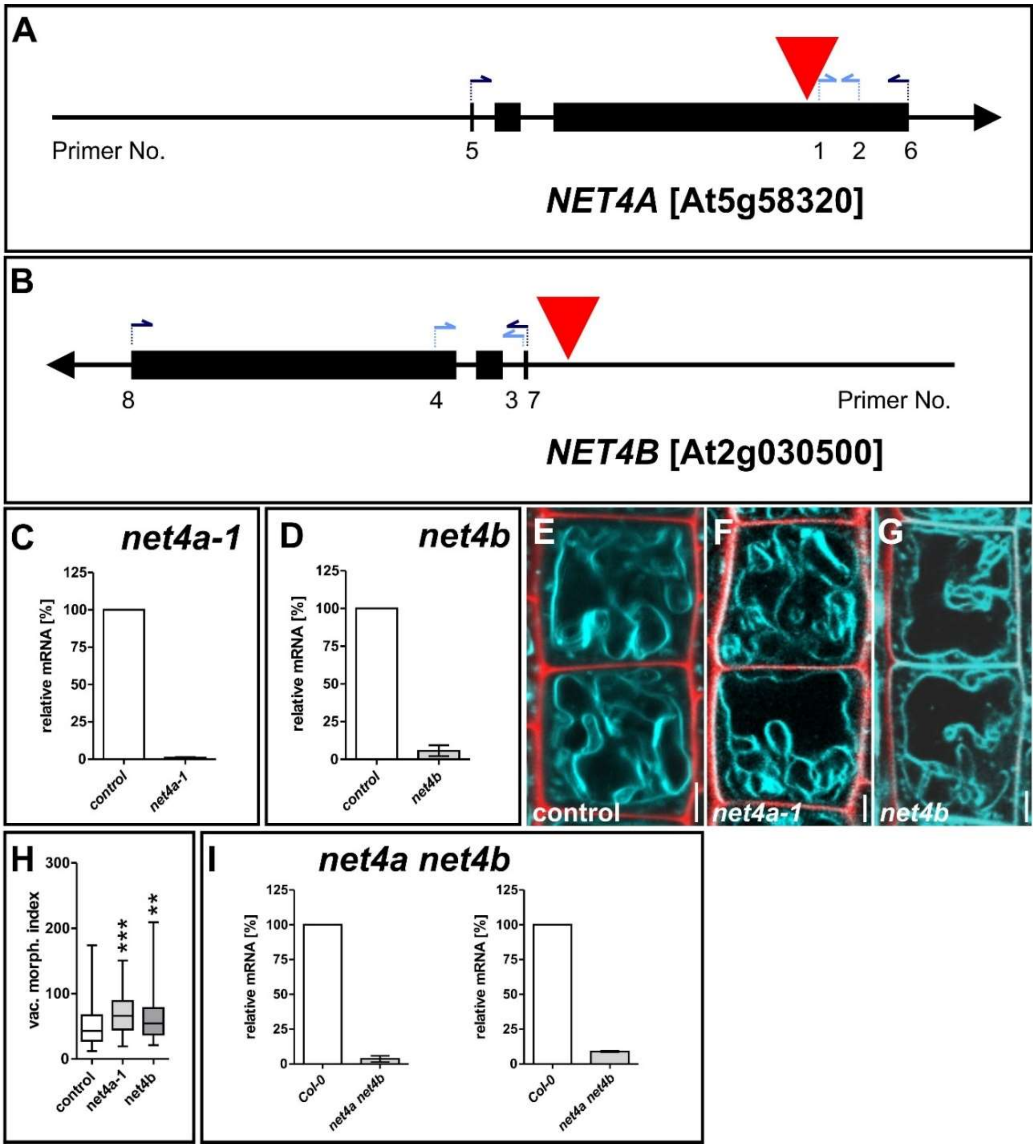


**Figure S2: Loss-of-function mutants of NET4A and NET4B.** Site of T-DNA insertion in the NET4A SALK line SALK_017623 **(A)** and NET4B line SALK_056957 **(B)**. qRT-PCR to test for gene expression levels **(C and D)**. Arrows indicate position of primers. **(E-G)** Vacuolar morphology of Col-0 control (n=126), the *net4a* (n=54) and the *net4b* single mutant (n=99). Quantification of vacuolar morphology **(H)**. After crossing of the single mutants, gene expression for *NET4A* and *NET4B* was tested again via qRT-PCR **(I)**. Primers used are listed in table S1. MDY-64 (cyan) was used to stain vacuoles, propidium iodide (red) to stain cell walls. Columns of bar charts represent mean values, error bars represent s.e.m. Box limits of boxblots represent 25th percentile and 75th percentile, horizontal line represents median. Whiskers display min. to max. values. Student's t-test, p-values: **p < 0.01; ***p < 0.001. Scale bars: 5 µm.
