## Supplementary material for "NET4 modulates the compactness of vacuoles in *Arabidopsis thaliana*": Figure S3

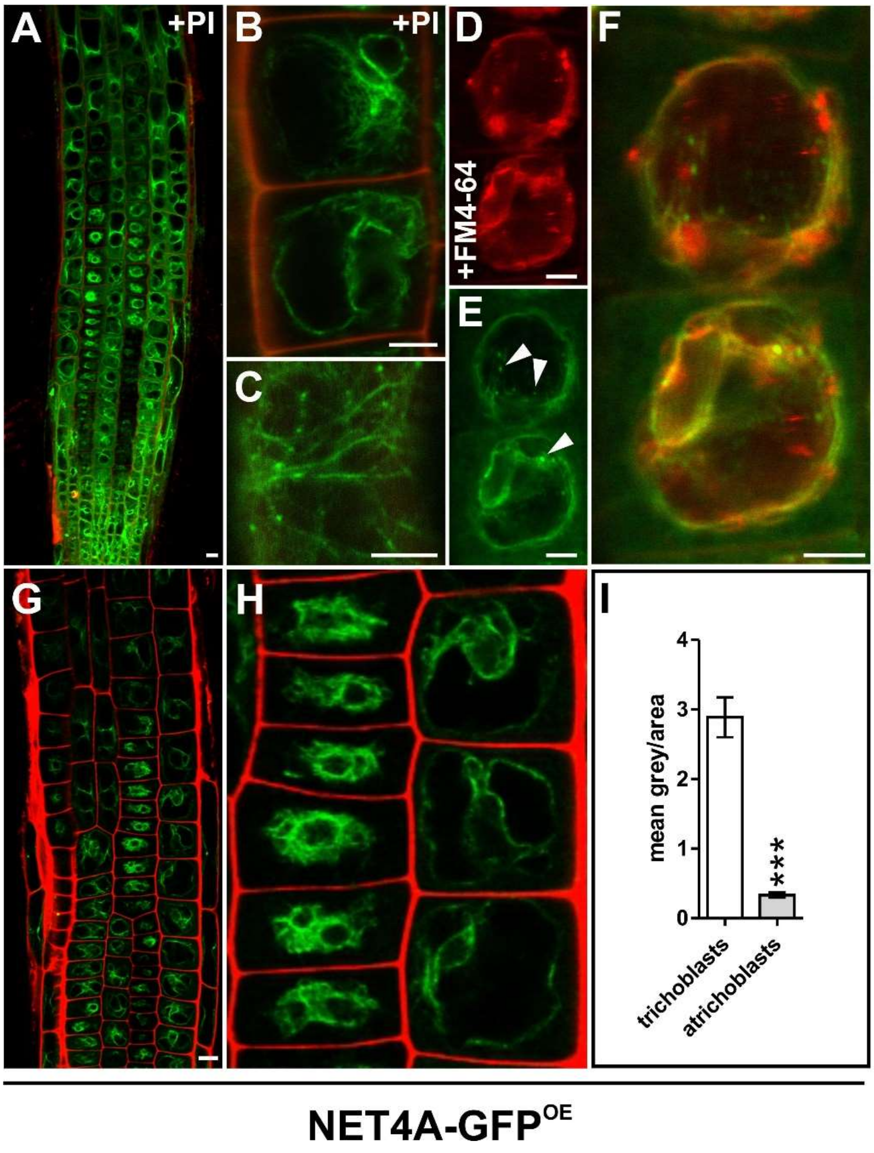


**Figure S3: Localization of NET4A driven by the CMV 35S promotor (NET4A-GFP$^{OE}$). (A)** Uniform signal distribution within the Arabidopsis root meristem. **(B)** Vacuolar signal in atrichoblast cells and **(C)** filamentous signal at the cell cortex. **(D-F)** Vacuole staining by FM4-64 (3 h) shows NET4A colocalization at the tonoplast. White arrowheads highlight punctate signals. **(G-I)** More constricted vacuoles in trichoblast cells (n=34) show a higher signal accumulation per area in comparison to less folded vacuoles in atrichoblast cells (n=35). Propidium iodide was used to stain cell walls, FM4-64 to stain the tonoplast. Error bars represent s.e.m. Student's t-test, p-values: ***p < 0.001. Scale bars: 5 µm.
