## Supplementary material for "NET4 modulates the compactness of vacuoles in *Arabidopsis thaliana*": Table S1

**Table S1: Used primers**

| Gateway Cloning Primers | Sequence |
|---|---|
| NET4A_FW | GGGGACAAGTTTGTACAAAAAAGCAGGCTTTATGGATTATGATCTGCTTCGTTC |
| NET4A_REV | GGGGACAAGTTTGTACAAAAAAGCAGGCTTAAGAAGCAAGAATGGATGATG |
| **Genotyping Primers** | **Sequence** |
| SALK_LBb1.3 | ATTTTGCCGATTTCGGAAC |
| SALK_017623(S1)_FW | AATGGATGATGGTCTTGTTGG |
| SALK_017623(S1)_REV | GAACACTGAGAGCTTGTTGCC |
| SALK_056957_net4b_FW | AATCACGATAGAGCCACATGC |
| SALK_056957_net4b_REV | TACATGCGGTAGAATTCCTCG |
| **qRT-PCR Primers** | **Sequence** |
| act2_fw2 | CTTGCACCAAGCAGCATGAAG |
| act2_rev | CCTGGACCTGCCTCATCATACTC |
| qRT_NET4A_fw (1) | GCTGGAAGCCAATGTGCGTTATC |
| qRT_NET4A_rev (2) | CGACTTATCTCCGACTCTAGCTCACTC |
| qRT_NET4B_fw (3) | CGTCTACGGCTCAGAGCAAG |
| qRT_NET4B_rev (4) | GCTGGATTAACTCGGGACGTTTC |
| UBQ-5_fw | GACGCTTCATCTCGTCC |
| UBQ-5_rev | GTAAACGTAGGTGAGTCCA |
| NET4A_FW (5) | GGGGACAAGTTTGTACAAAAAAGCAGGCTTTATGGATTATGATCTGCTTCGTTC |
| NET4A_REV (6) | GGGGACAAGTTTGTACAAAAAAGCAGGCTTAAGAAGCAAGAATGGATGATG |
| NET4B_FW (7) | GGGGACAAGTTTGTACAAAAAAGCAGGCTTTATGGCTTCGTCTACGGCTCAG |
| NET4B_REV (8) | GGGGACAAGTTTGTACAAAAAAGCAGGCTTTCAAGTTGATAAGACCACATACTCTCTT |
